# Identifying disease-associated dysbiosis in image-based spatial microbiome via MiSA

**DOI:** 10.64898/2026.11.16.767125

**Authors:** Ruoqian Liu, Benjamin Grodner, Sarah Samorodnitsky, Daxuan Deng, Asmita Roy, Yuzheng Dun, Boyi Guo, Iwijn De Vlaminck, Ni Zhao, Wodan Ling

**Affiliations:** Division of Biostatistics, Department of Population Health Sciences, Weill Cornell Medicine, 425 E 61st Street 10128, NY, USA; School of Oceanography, University of Washington, 616 NE Northlake Place, 98105, WA, USA; Department of Epidemiology and Biostatistics, Memorial Sloan Kettering Cancer Center, 485 Lexington Avenue, 10017, NY, USA; Department of Biostatistics, Johns Hopkins Bloomberg School of Public Health, 615 N Wolfe Street, 21205, MD, USA; Public Health Sciences Division, Fred Hutchinson Cancer Center, 1100 Fairview Ave N, 98109, WA, USA; Division of Biostatistics, Department of Population Health Sciences, University of Utah, 295 Chipeta Way, 84108, UT, USA; Nancy E and Peter C Meinig School of Biomedical Engineering, Cornell University, 237 Tower Road, 14853, NY, USA

**Keywords:** Image-based spatial microbiome, Spatial dysbiosis, Deviation from complete spatial randomness, Omnibus test, Multiscale modeling

## Abstract

**Background:** Extensive studies have revealed the associations of microbiome dysbiosis with disease through compositional analysis of 16S rRNA amplicon or shotgun metagenomic sequencing data. However, dysbiosis in spatial organization of microbiome inhabiting the body sites and its implication for diseases remain understudied. Recently developed high phylogenetic resolution fluorescence in situ hybridization (HiPR-FISH) is an image-based mapping technology that provides spatial microbiome data at single-cell resolution, which allows for visualizing and quantifying spatial arrangement of microbial cells. Despite the growing availability of such data, no methods have been tailored to studying spatial dysbiosis in image-based microbiome data.

**Results:** Here we introduce MiSA (Microbiome Spatial Aberration), the first statistical framework for identifying disease-associated spatial dysbiosis and the individual taxa that may contribute to it, based on differing degrees of clustering or regularity, i.e., microbial cells being more aggregated or more evenly spaced than expected under complete spatial randomness, between diagnostic groups. In simulations based on a real dataset, MiSA was robust across diverse and noisy spatial configurations, achieving substantially higher statistical power than a method borrowed from spatial proteomics, while controlling type I error at both the individual-taxon and community levels. When applied to an oral biofilm study, MiSA detected peri-implant mucositis-associated spatial dysbiosis at the community level and identified five taxa as potential contributors, including *Actinomyces*, which showed increased clustering in mucositis.

**Conclusions:** MiSA is a powerful and robust framework for identifying spatial dysbiosis and the contributing taxa in image-based microbiome data, offering new insights into the spatial organization of microbial communities in disease.

## 1 Background

Microbial balance in the human body is closely related to individual health [1]. Disruption of the microbial ecosystem can lead to an imbalance between beneficial and pathogenic bacteria. This imbalance, known as dysbiosis, refers to a deviation from the usual microbial composition observed in healthy hosts and has been implicated in a wide range of diseases [2–4].

Although microbiome dysbiosis has been extensively investigated through compositional analysis of 16S rRNA amplicon or shotgun metagenomic sequencing data [4], alterations in microbial spatial organization remain understudied. Recently developed high phylogenetic resolution fluorescence in situ hybridization (HiPR-FISH) [5], an image-based microbiome mapping technology, has been applied to the mouse gut microbiome and created bacterial maps across multiple tissue sections. This approach enables exploration of microbial biogeography at single-cell resolution, making it possible to visualize and quantify the spatial arrangement of bacterial cells. A recent study applied HiPR-FISH to oral biofilms collected around dental implants from individuals with clinical diagnosis, including peri-implant health, peri-implant mucositis, and peri-implantitis [6]. The study targeted 18 prevalent microbial genera and generated spatial maps for the microbial cells. Figure 1 shows representative images from peri-implant health (a,b) and peri-implant mucositis (c,d). Rather than a homogeneous mixture of different species randomly distributed on the colonized surface, microbes within oral biofilms are arranged nonrandomly, forming spatially structured polymicrobial communities [7–9]. With such high-resolution spatial data, it is essential to assess whether disease-associated spatial dysbiosis occurs within the microbial community and to identify the taxa that contribute to the shifts in spatial organization.

**Fig. 1.**
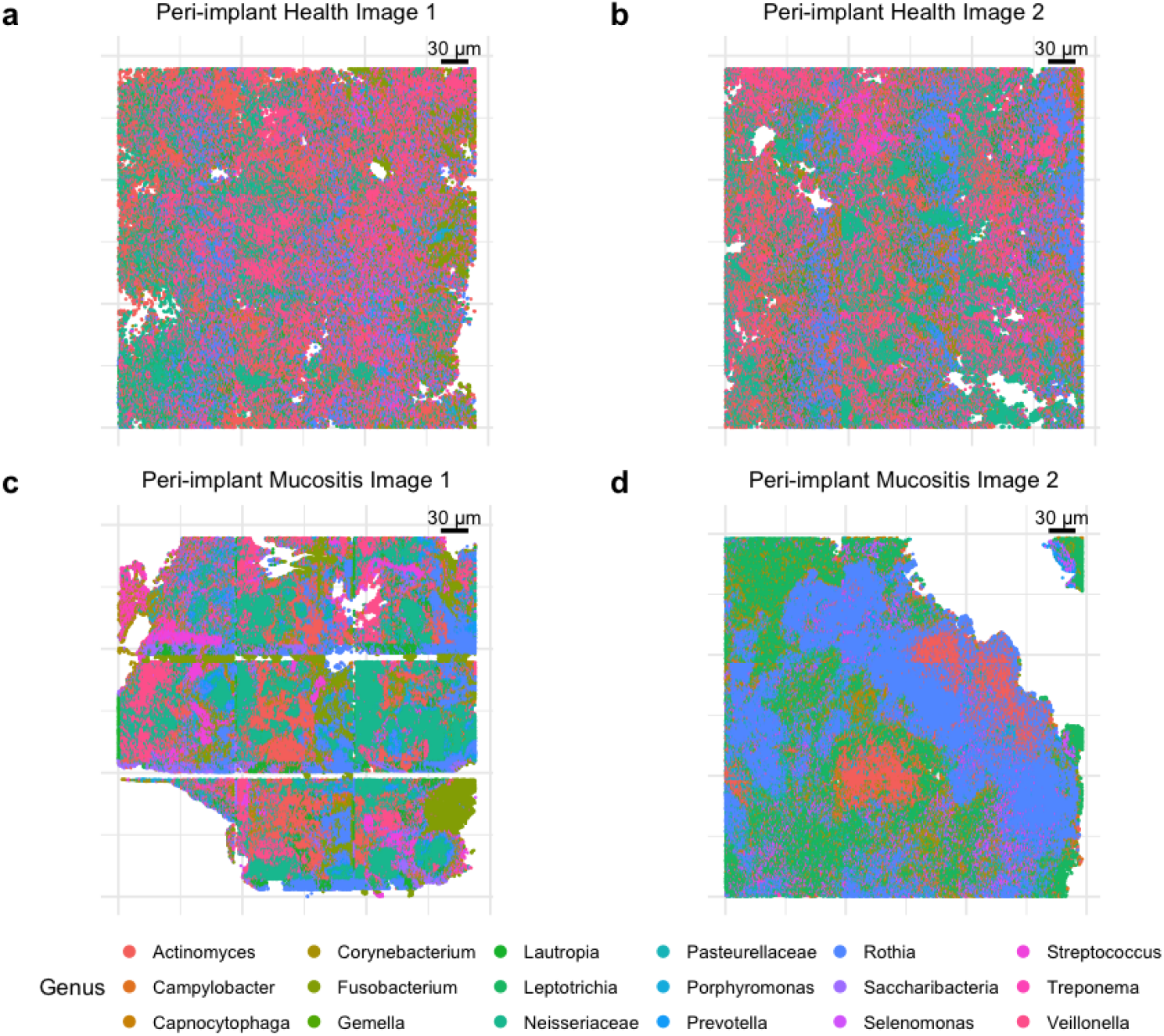
Representative examples of segmented oral biofilm spatial microbiome data with genus identities.

Despite the growing availability of such data, no methods have been tailored to image-based spatial microbiome data, which record the locations and taxonomic identities of individual microbial cells and thus form a spatial configuration of multiple cell types. Methods borrowed from spatial transcriptomics or proteomics to characterize cell-type spatial distributions are insufficient when applied directly, because they do not address challenges specific to these data.

First, spatial microbiome data typically include tens of taxa (depending on the taxonomic level) and thus form a multi-type point pattern. Methods developed for other spatial omics data largely focus on pairwise co-occurrence or interactions among a handful of selected cell types. For example, spicyR [10] and SpaceANOVA [11] use spatial summary functions such as Ripley’s *K*, Besag’s *L*, and the pair correlation function *g*, whereas DIMPLE [12] compares kernel-smoothed intensities between pairs of cell types. When applied to the entire microbial community, the number of pairs grows quadratically with the number of taxa (e.g., 153 pairs for the 18 taxa in our motivating study), producing many heterogeneous pairwise results and a heavy multiple-testing burden that reduces statistical power. Spatial entropy-based methods, an alternative direction, consider all cell types simultaneously. For example, FunSpace [13] applies functional principal component analysis (FPCA) to relate spatial entropy curves to survival outcomes. However, entropy summarizes the diversity of cell types and depends only on their proportions, not on their identities, thus, regions with different taxon compositions can yield identical entropy values (e.g., a region in which taxa A and B are evenly intermixed has the same entropy as a region in which taxa C and D are evenly intermixed), so differences in the spatial arrangement of individual taxa may be obscured.

Second, spatial microbiome data show substantial heterogeneity even within diagnostic groups. Functional data analysis (fda)-based methods that link spatial summary functions to clinical outcomes [14] rely primarily on systematic shifts in the functional trajectories. When spatial arrangements vary widely within groups, such shifts are weak relative to the within-group variability, resulting in reduced statistical power.

Third, bacteria naturally form colonies, so each taxon exhibits clustering at small spatial scales even in well-mixed communities. Methods that test each image against complete spatial randomness (CSR) will therefore detect colony formation itself, masking the between-group differences of interest. Moreover, microbial spatial signals often vary with spatial scale. A taxon may be clustered at short distances, reflecting the aggregation of cells within colonies, but appear regular at longer distances, where its spatial summary function instead reflects how the colonies themselves are spaced across the image. As a result, its deviations from randomness occur in opposite directions at different scales. Methods that aggregate these signed deviations across distances into a single statistic, such as spicyR, may cancel out these opposing signals.

To address these limitations, we propose MiSA (Microbiome Spatial Aberration), the first hypothesis testing framework that effectively identifies spatial alterations of the microbiome with respect to disease. Within this framework, each taxon is evaluated individually to identify differences in its spatial pattern between the disease and healthy groups. Specifically, we propose a spatial summary function *d* that quantifies each point pattern’s deviation from CSR across spatial scales, and compare these deviations across groups rather than testing each image against CSR, so that MiSA is not confounded by clustering shared by all images, such as colony formation. MiSA’s multiscale comparisons capture group differences even when spatial patterns vary widely within groups, without relying on systematic shifts of entire curves. Spatial dysbiosis can be further assessed by combining these taxon-specific results to generate an overall inference for the microbial community.

MiSA demonstrates high sensitivity and robustness in detecting spatial dysbiosis in various spatial configurations between diagnostic groups. In simulation studies, MiSA controlled the type I error and achieved substantially higher power than a competing method borrowed from spatial proteomics. Applied to the oral biofilm study [6], MiSA identified spatial dysbiosis between peri-implant health and peri-implant mucositis groups and the contributing genera, providing novel biological insights into disease progression around dental implants.

## 2 Methods

### 2.1 Quantifying deviation from CSR with the d curve

Following segmentation of microbial cells and taxonomic classification, each cell is represented as a point in two-dimensional space, defined by its spatial coordinates and taxonomic identity (Figure 1). Each image thus becomes a microbial map, a multi-type point pattern that can be naturally modeled as point processes.

Considering one taxon in the spatial microbiome context, various spatial point patterns may be present. For example, if the microbial cells are independently and uniformly distributed over the space, the observed points follow a homogenous Poisson process, also known as complete spatial randomness (CSR). In most other cases, microbial cells can be clustered, regular (also called dispersed or ordered, where points are spread out evenly, e.g., in a grid or hexagonal pattern), or a combination of these spatial patterns. These characteristics of point patterns can be summarized by Ripley’s *K* function [15, 16] which counts the number of points within distance *r* of a randomly chosen point. Ripley’s *K* is defined as:

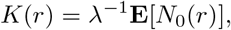

where **E** is the expectation operator, *N*_0_(*r*) is the number of points found within radius *r* of point *x*_0_ (an arbitrary point of the point pattern), and *λ* is a constant parameter describing the intensity of points in the image, implying that the expected density of microbial cells does not vary across spatial locations. Under CSR, *K*(*r*) = *πr*^2^ for all radii *r*. If the observed *K*(*r*) exceeds its theoretical value under CSR, this suggests spatial clustering of points, and the reverse indicates a regular spatial point pattern. The *K* function can be transformed to the variance-stabilized *L* function by

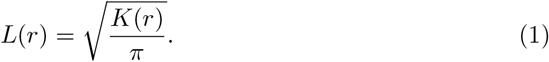

Similarly, under CSR, it can be shown that *L*(*r*) = *r* for all *r*, making it more convenient to interpret and visualize than the *K* function.

To quantify the deviation of the *L* curve from its theoretical values under CSR at any radius *r*, we propose a summary function

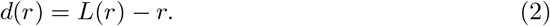

Notably, *d*(*r*) = 0 under CSR. When *d*(*r*) *>* 0, clustering is indicated, whereas *d*(*r*) *<* 0 suggests regularity. The magnitude reflects the strength of the departure from CSR. In practice, point patterns of spatial omics data often exhibit a combination of these characteristics across different spatial scales, i.e., as *r* varies.

In practical applications with observed data, *K*(*r*) can be empirically estimated as

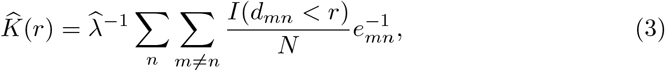

where *λ* is estimated as 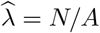, with *N* being the total number of points and *A* being the area of the study region, *d*_*mn*_ is the distance between the *m*th and *n*th points, *I*(*·*) is the indicator function, and *e*_*mn*_ is the edge correction factor that accounts for image boundaries (commonly used edge correction methods are reviewed by Baddeley [17]). The estimate 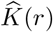 is essentially the average number of microbial cells located within a radius *r* of a given cell. We can then obtain estimates, 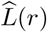 and 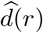, following formulas (1) and (2).

To illustrate how *d*(*r*) characterizes the spatial arrangement of point events, we consider two point patterns shown in Figure 2(a, b), generated over the unit square. Point Pattern 1 represents a microbial species that forms small colonies scattered randomly throughout the image. In contrast, Point Pattern 2 represents the spatial expansion of microbial colonies, resulting in fewer but larger clusters, and thus exhibits a stronger clustering pattern. Both point patterns are realizations of a Matérn cluster process [18], generated with the `rMatClust` function in the R package `spatstat`, with a cluster radius of 0.03 for Point Pattern 1 and 0.1 for Point Pattern 2 (see Additional file 1: Section S1 for details).

**Fig. 2.**
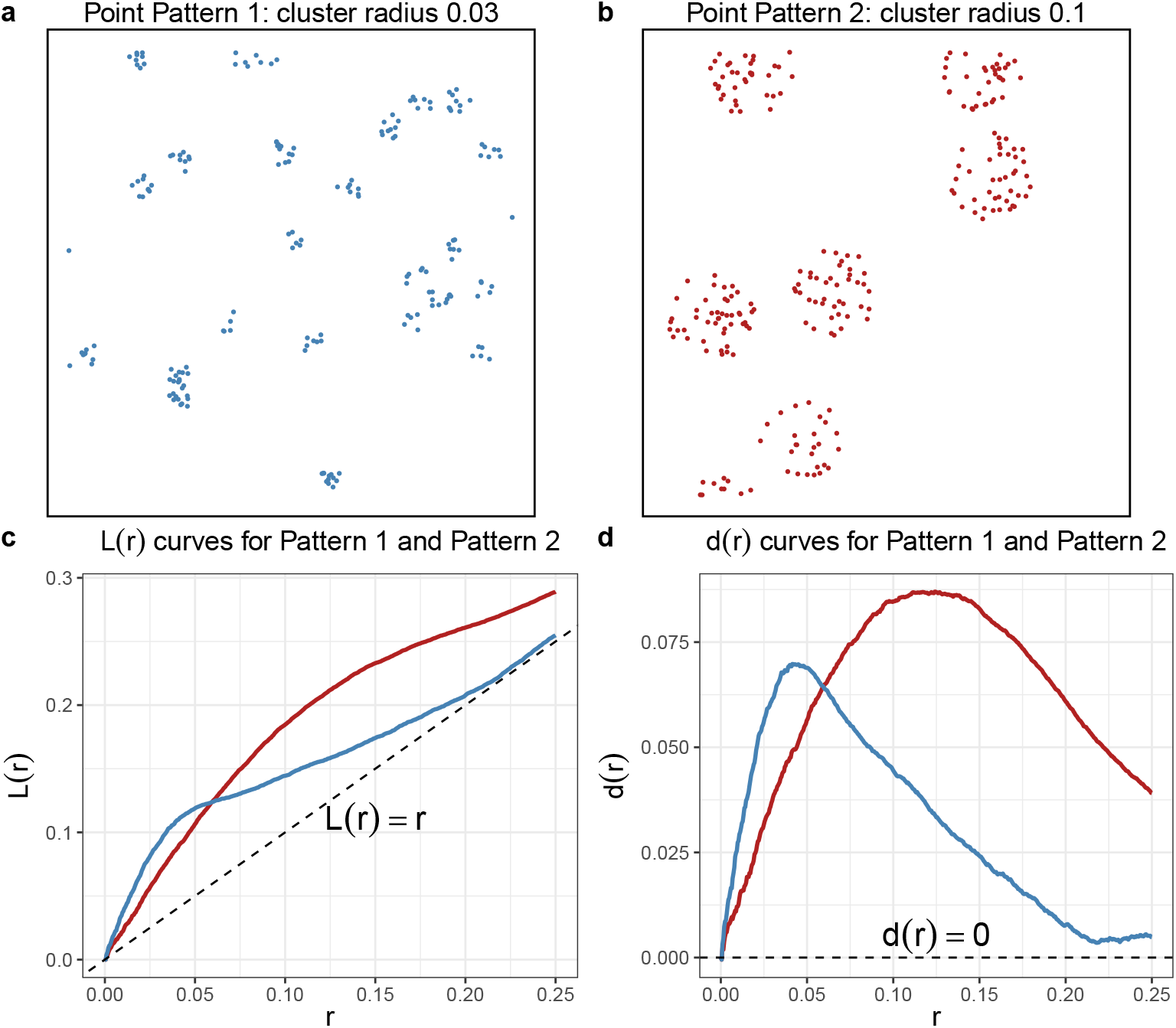
(a) The Matern’s cluster pattern with cluster center intensity (kappa) 25 and radius 0.03. The number of points per cluster follows a Poisson distribution with mean mu = 8. (b) The Matern’s cluster pattern with kappa = 5 and radius 0.1, and mu = 40. Both patterns have the same overall intensity kappa * mu = 200. (c, d) *L* and *d* curves as functions of distance *r* ∈ [0, 0.25] for the two patterns, with dashed lines representing CSR.

Figure 2 shows the estimated *L* and 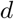 curves for the two patterns. Over *r* ∈ [0, 0.25], 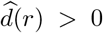 for both patterns, indicating clustering across these spatial scales, with larger values corresponding to stronger clustering. At short distances (approximately *r* ≤ 0.06), Pattern 1 has larger 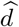 values (i.e., more neighboring points) than Pattern 2, whereas Pattern 2 has larger values for *r* ∈ [0.06, 0.25]. This is intuitive, as the tighter clusters in Pattern 1 produce an excess of nearby points at short distances, whereas the more dispersed clusters in Pattern 2 produce an excess of points at larger distances.

Beyond the scale-specific comparison, the peak absolute magnitude of 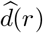, which can occur at any distance *r*, represents the maximum deviation from CSR over the entire range of *r* considered. In this example, both peaks are positive, and Pattern 2 shows a higher peak than Pattern 1, indicating a larger maximum deviation from CSR and thus stronger clustering. Alternatively, the area between the 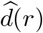 curve and the CSR baseline integrates absolute deviations across all distances. This area is also larger for Pattern 2, indicating stronger clustering overall. These two summaries, the peak and the integrated deviation, allow spatial patterns to be compared globally.

This example demonstrates that the summary function *d*(*r*) effectively captures spatial signals that vary across scales.

### 2.2 Detecting taxon-specific differences in spatial patterns

MiSA (Figure 3) leverages *d*(*r*) to differentiate diagnostic groups based on taxon-specific spatial patterns and to assess microbiome spatial dysbiosis associated with disease. For taxon *g*, let 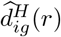 denote the estimated *d* curve for image *i* in the Healthy group, and 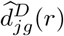 for image *j* in the Disease group. These curves are evaluated over a set of radii, e.g., *r* = 6, 12, 18, …, *r*_max_, where *r*_max_ is the maximum radius, determined based on the specific data and scientific question. Since each microbial taxon *g* is analyzed individually, the subscript *g* is omitted hereafter for simplicity.

**Fig. 3.**
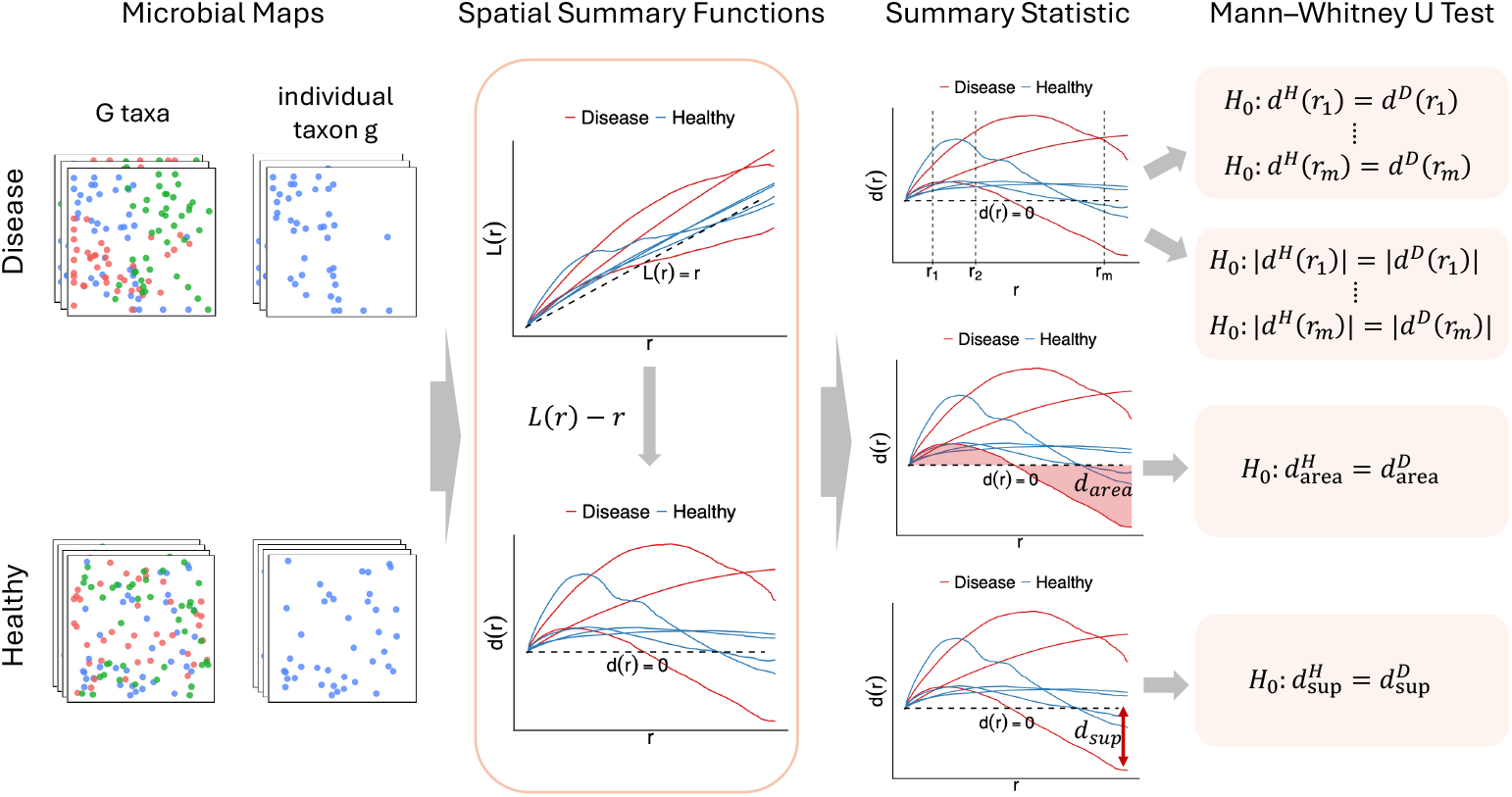
Overview of the MiSA workflow. Image-based spatial microbiome data for each taxon *g* are analyzed individually using the summary function *d*. Four test statistics are derived from *d* to detect differences in spatial patterns between the two groups. Taxon-specific *p*-values, *p*_*g*_, are obtained using Mann–Whitney U test, and are combined using the HMP method to yield an inference for the overall dysbiosis of the microbial community.

#### MiSA-orig and MiSA-abs

Detecting spatial differences between groups at multiple specific radii is informative for two reasons. First, for some taxa, the differences between diagnostic groups arise only at particular spatial scales, appearing at those scales and vanishing at others. Second, *d*(*r*) can be positive or negative and may change sign across radii. We therefore propose to compare deviations from CSR in two ways. First, comparing the raw statistic *d*(*r*) accounts for both the magnitude and the direction of deviation (clustering or regularity), testing whether the excess or deficit of neighbors at a given radius differs between groups. Alternatively, comparing |*d*(*r*)| considers only the magnitude of deviation, testing whether the groups differ in how far their spatial patterns depart from randomness, with clustering and regularity treated equivalently. Thus, the global null hypotheses are

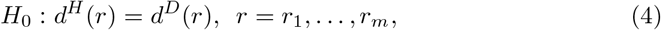

and

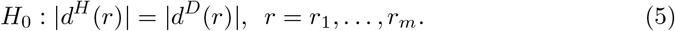

For tests (4) and (5), we apply the Mann–Whitney U test at each pre-selected radius, yielding *p*-values 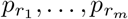. To aggregate signals across scales, we combine these *p*-values using the harmonic mean *p*-value (HMP) method [19], a widely adopted approach for combining dependent *p*-values while controlling the family-wise error rate (FWER). The combined *p*-value serves as the taxon-level test result.

#### MiSA-sup and MiSA-area

In addition to testing at specific radii, we can assess the spatial structure of each image as a whole by extracting image-level summary statistics that capture the spatial signal across radii. As illustrated in Figure 2, one option is to compare the peak signal between diagnostic groups, which reflects the most striking spatial pattern in an image, summarized by the maximum absolute deviation from CSR, 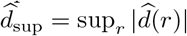. Alternatively, one can compare the integrated deviation, which reflects the overall strength of the spatial signal, summarized by the area between 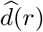 and the CSR baseline, 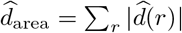.

We take the absolute value of 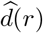 in both summaries so that deviations from CSR in either direction contribute to the statistics equally. For example, for a taxon that is clustered at short distances 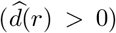 and regularly spaced at longer distances 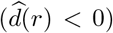, integrating 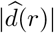 accumulates both deviations rather than letting them cancel out. Based on 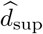 and 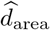, we test

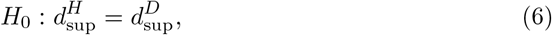

and

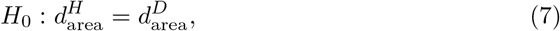

using Mann–Whitney U test and obtaining *p*-values.

#### Effect sizes

We can calculate the effect size for MiSA-sup and MiSA-area as 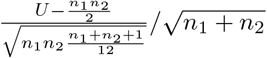[20], where *U* is the Mann–Whitney U statistic, and *n*_1_ and *n*_2_ are the sample sizes of the two groups. A positive value indicates that the reference group tends to have a larger deviation from CSR than the comparison group, while a negative value indicates the reverse. The absolute value of the effect size is commonly classified as small (≥0.1), medium (≥0.3), or large (≥0.5). Note that effect sizes are not available for MiSA-orig and MiSA-abs, as these are omnibus tests that combine *p*-values across radii. Their single summary *p*-values reflecting overall significance rather than a signed statistic from which an effect size could be derived.

### 2.3 Assessing spatial dysbiosis of microbial community

In hypothesis tests (4) - (7), each taxon is analyzed individually, yielding a taxon-specific *p*-value *p*_*g*_. We assess the overall microbiome spatial dysbiosis by aggregating results for all taxa of interest, i.e., combining the individual *p*_*g*_ for *g* = 1, …, *G*, using the HMP method. If the HMP combined *p*-value is less than the pre-set significance level, we conclude that the microbial profile exhibits spatial dysbiosis that is associated with disease. Taxon-level *p*-values are interpreted as exploratory, to identify taxa that may contribute to the community-level signal and to guide further investigation. When there is no specified biological hypothesis, MiSA-sup is the default, as it does not assume that signals occur at particular radii and is sensitive to strong signals at any spatial scale.

### 2.4 Numerical Experiments and Simulations

To assess MiSA in terms of type I error and statistical power, we conducted simulation studies under a range of settings. For all simulations, image-based spatial microbiome data were generated using templates derived from real peri-implant biofilm images [6]. Each image profiled 18 microbial genera, with spatial locations recorded for segmented microbial cells.

Because no tools have been developed for analyzing spatial dysbiosis of image-based microbiome data, we compared MiSA with a functional data analysis (fda) approach developed for spatial proteomics data [14]. This approach treats spatial summary functions, such as Ripley’s *K*, Besag’s *L*, or spatial entropy, as functional data and tests whether they are associated with a clinical outcome. To keep the comparison consistent with MiSA, we used the estimated *L* function of each taxon as the functional predictor and fitted a scalar-on-function regression with diagnostic group as the binary outcome, using the `pfr` function in the R package `refund`. Similar to MiSA, the competing test was performed at the taxon level to identify spatial differences, and the taxon-specific *p*-values were then combined to obtain inference at the community level. We refer to this competing method as “fda” hereafter.

#### 2.4.1 Type I error simulations

To generate data under the null hypothesis, we randomly sampled 40 images from real biofilm images and randomly assigned them to two groups of equal size (20 images per group). To preserve the original spatial organization of the microbiome while introducing variability across simulation replicates, we randomly subsampled 60% of the microbial cells from original images in each replication.

We applied MiSA and fda to the simulated dataset, obtaining taxon-specific *p*-values *p*_*g*_ for *g* = 1, 2, …, 18. These *p*-values were then combined using HMP to yield a *p*-value for inferring overall microbiome spatial dysbiosis. We repeated the simulation 1000 times and summarized the empirical type I errors at both the individual-taxon and microbial community levels as the proportion of *p*-values less than *α* = 0.05.

#### 2.4.2 Power simulations

To assess power for identifying spatial dysbiosis associated with disease, we generated datasets following a similar procedure under the alternative hypothesis. 40 null images were generated by subsampling 60% of microbial cells from real images, and were then randomly assigned to two groups of equal size. For the healthy group, spatial distributions of all 18 genera remain unchanged.

For the disease group, spatial alterations were introduced for five selected genera (*Gemella, Porphyromonas, Prevotella, Treponema*, and *Pasteurellaceae*). Differential spatial patterns were created by superimposing additional microbial cell clusters onto the null images. These clusters were generated using a Thomas cluster process implemented via the `rThomas` function from the `spatstat` R package, with cluster center intensity `kappa = 1e-7` (controlling the number of additional clusters on the image) and cluster size `scale = 300`. Effect size was varied by adjusting the mean number of points per cluster, `mu` ={250, 350, 550}, with larger values corresponding to denser clusters and thus stronger spatial patterns.

Both taxon-level and community-level *p*-values were obtained by applying MiSA and fda to the simulated dataset. Statistical power was estimated as the proportion of *p*-values less than *α* = 0.05 across 1000 simulation runs.

### 2.5 Application to the oral biofilm dataset

#### 2.5.1 Data overview

Oral biofilms were collected from donors diagnosed with peri-implant health, peri-implant mucositis and peri-implantitis, and processed by HiPR-FISH [6]. The study targeted 18 abundant and prevalent microbial genera and imaged the stained samples using confocal microscopy. Using automated image segmentation, microbial cells were classified based on their fluorescent spectra, after which spatial maps of genus identity were generated (example images are shown in Figure 1). We focused on 14 images from healthy individuals and 19 images from peri-implant mucositis patients to identify differences in spatial patterns between the two clinical groups.

#### 2.5.2 Determine the true region of study

In the oral biofilm study, bacterial maps often contain blank regions where no microbial cells are observed. These empty areas can introduce additional inhomogeneity and lead to spatial patterns of microbial taxa that are merely artifacts rather than true biological structure. Therefore, accurately specifying the region of study ensures valid estimation of spatial summary statistics (the *d* curves) and reliable hypothesis testing results by MiSA.

We presented a preprocessing step to delineate the true study region, as illustrated in Figure 4. Specifically, we considered microbial cells of all genera in each image as a spatial point process (Figure 4a). We then performed kernel density estimation for local point intensity. The smoothing bandwidth was selected via cross-validation. The estimated density map is shown in Figure 4b. Regions with kernel density below 0.002 were identified as outliers relative to the main distribution of region density (Figure 4c) and were therefore excluded from further analysis. Because the imaging protocol produced images of consistent quality, a threshold of 0.002 served as a clear cutoff and was applied to all biofilm images. In addition, small isolated “islands” separated from the main microbiome-colonized area were removed. To accommodate the irregular geometry and non-convex shape of the remaining region, we applied the *α*-convex hull [21] to define the true region of study for each image. The resulting study region (Figure 4d) was used in the subsequent tests.

**Fig. 4.**
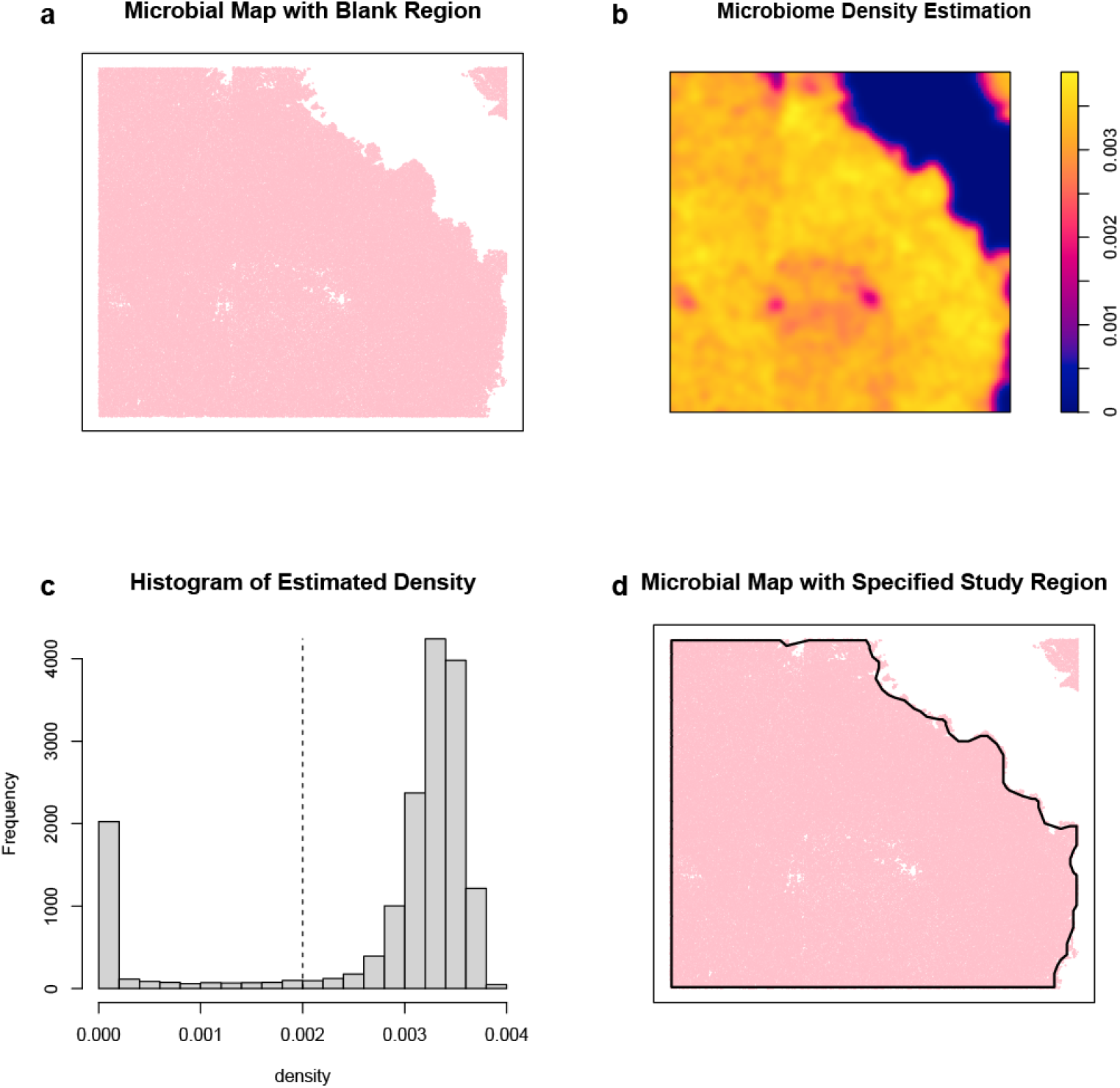
(a) Bacterial maps with blank regions where no microbial cells are observed. (b) Kernel density estimation for local point intensity. (c) Histogram of estimated density. Regions with kernel density below 0.002 were excluded from analysis. (d) Apply the *α*-convex hull to determine the study region.

## 3 Results

### 3.1 Simulation results

At the taxon level (Figure 5a), type I errors for MiSA-orig and MiSA-abs were consistently below 0.05 for all 18 taxa, while MiSA-sup and MiSA-area ranged from 0.033 to 0.057. In contrast, the fda method is conservative, with type I error below 0.016 for all 18 taxa. At the community level (Figure 5b), type I errors for the four tests of MiSA range from 0.041 to 0.057. These findings confirm that MiSA is a valid method that controls type I error at both the individual-taxon and community levels.

**Fig. 5.**
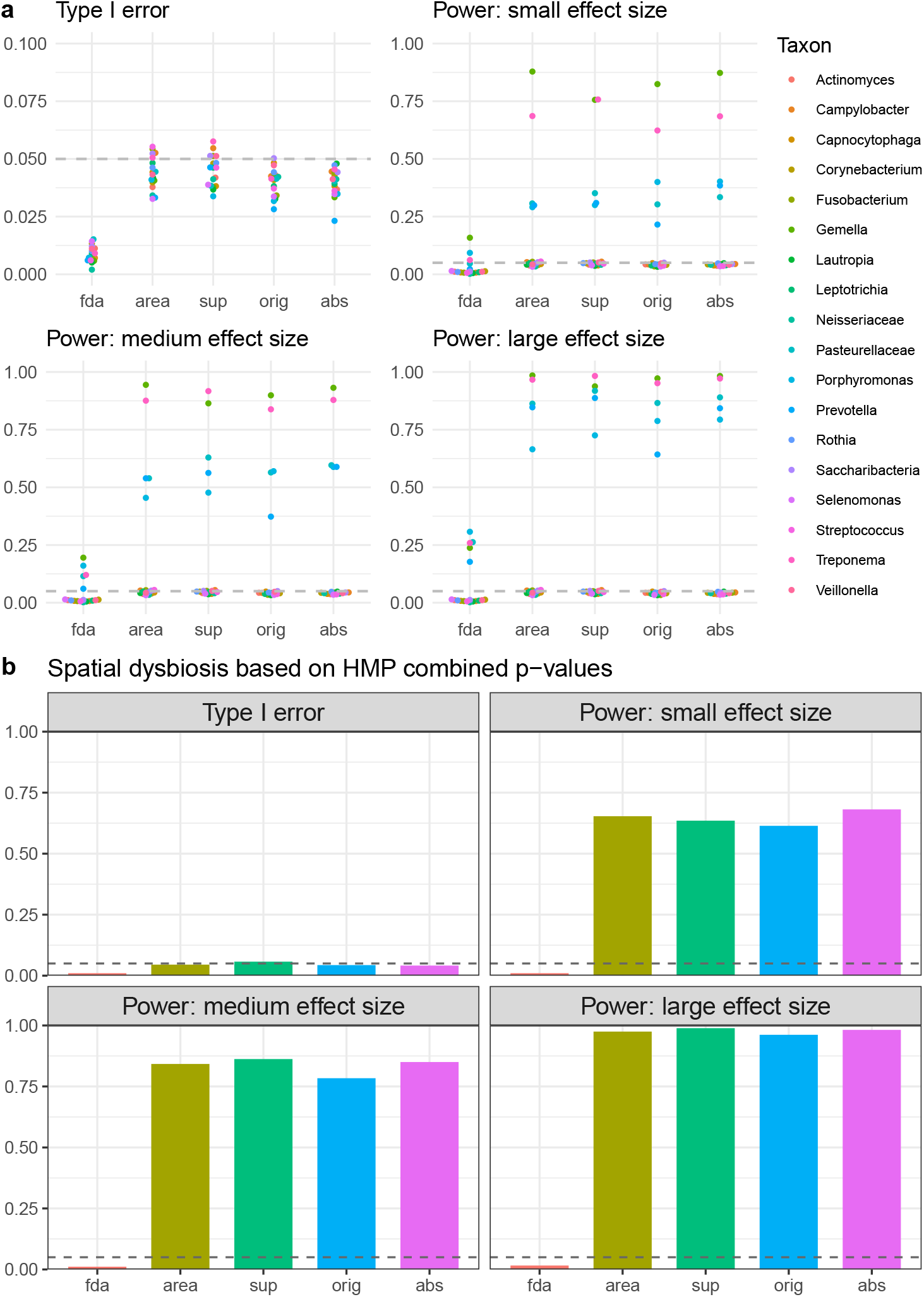
Simulation results for MiSA and the competing method. (a) type I error and power (small, medium and large effect size) based on taxon-specific *p*-values. (b) type I error and power (small, medium and large effect size) based on combined *p* -values across all taxa. The four tests of MiSA are area, sup, orig, and abs, while fda is the competing method.

Taxon-level power was evaluated as the rejection rate for the five taxa, *Gemella, Porphyromonas, Prevotella, Treponema*, and *Pasteurellaceae*, with true differences in spatial pattern (Figure 5a). The power of both methods increased with effect size, but MiSA achieved substantially higher power than fda for all five taxa at every effect size. In particular, fda had limited ability to detect weaker signals. At the small effect size (`mu = 250`), its power was below 0.05 for *Prevotella* and *Pasteurellaceae*, and it exceeded 0.05 for all five taxa only at the large effect size (`mu = 550`). For the remaining 13 taxa without spatial differences, MiSA’s rejection rates remained around 0.05, further confirming type I error control at the taxon level. We assessed statistical power for detecting spatial dysbiosis based on the HMP combined *p*-value across the 18 taxa. The results in Figure 5b demonstrate that MiSA outperforms the competing method at the community level, maintaining superior power for varying effect sizes.

### 3.2 Oral biofilm data analysis

The radius *r* ranged from 0 to 3000 pixels (1 pixel *≈* 0.0675 *µ*m) in steps of 6 pixels, i.e., *r* = 0, 6, 12, …, 3000. For each taxon, we excluded images with limited spatial range, and truncated the remaining curves to a common maximum *r*. Figure 6 shows the 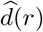 curves for the 18 microbial genera, with each curve representing one image and colored by its diagnostic group. For MiSA-orig and MiSA-abs, we evaluated 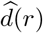 and 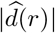 at 20 pre-specified radii evenly spaced over the range of *r*. For MiSA-sup and MiSA-area, the summary statistics 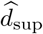 and 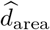 were computed from 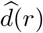 evaluated over all radii. The *p*-values and effect sizes (with the healthy group as the reference) are reported in Figure 7 and Tables 1 and 2.

**Table 1.**
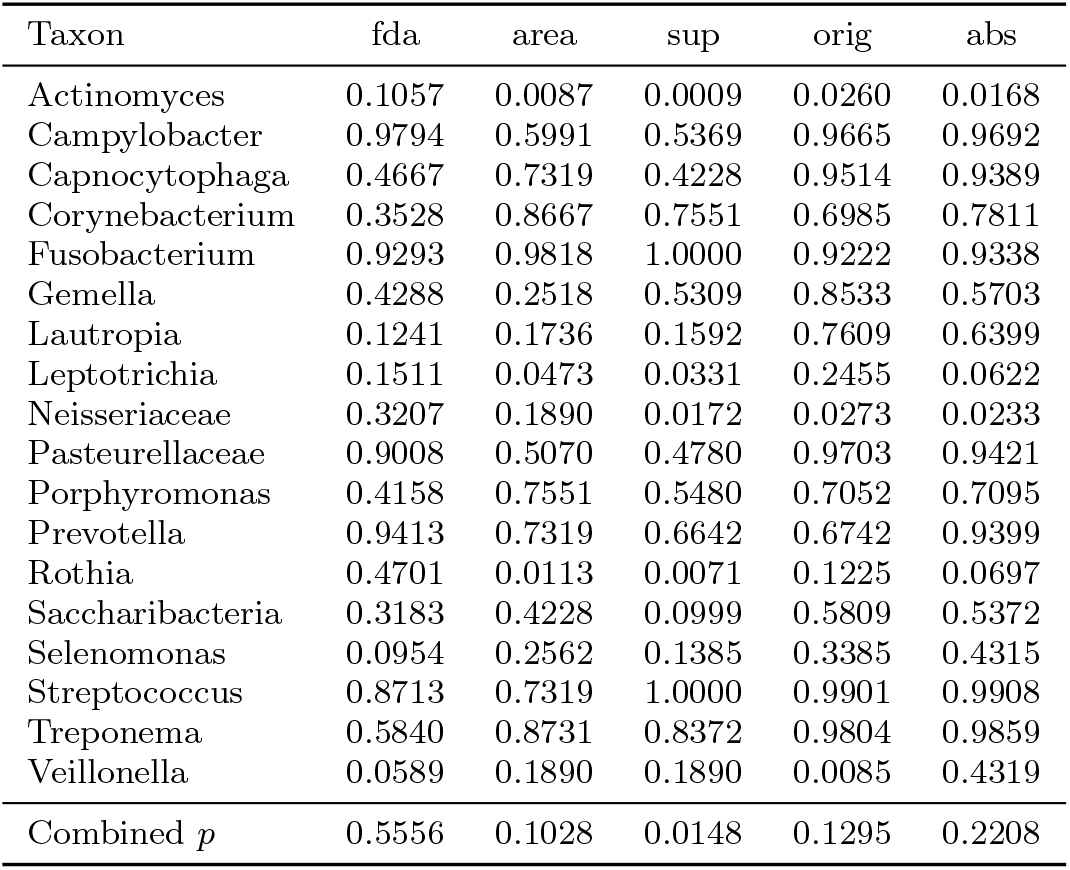
*p*-values for Oral Biofilm Analysis.

**Table 2.** Effect sizes for Oral Biofilm Analysis.

| Taxon | area | sup |
|---|---|---|
| Actinomyces | -0.498 | -0.610 |
| Campylobacter | 0.105 | 0.123 |
| Capnocytophaga | -0.070 | -0.158 |
| Corynebacterium | -0.038 | -0.066 |
| Fusobacterium | 0.009 | 0.000 |
| Gemella | -0.242 | -0.136 |
| Lautropia | -0.263 | -0.272 |
| Leptotrichia | 0.377 | 0.404 |
| Neisseriaceae | -0.254 | -0.447 |
| Pasteurellaceae | -0.132 | -0.140 |
| Porphyromonas | -0.066 | -0.122 |
| Prevotella | -0.070 | -0.088 |
| Rothia | -0.474 | -0.500 |
| Saccharibacteria | -0.158 | -0.316 |
| Selenomonas | -0.225 | -0.291 |
| Streptococcus | 0.070 | 0.000 |
| Treponema | -0.035 | -0.044 |
| Veillonella | -0.254 | -0.254 |

**Fig. 6.**
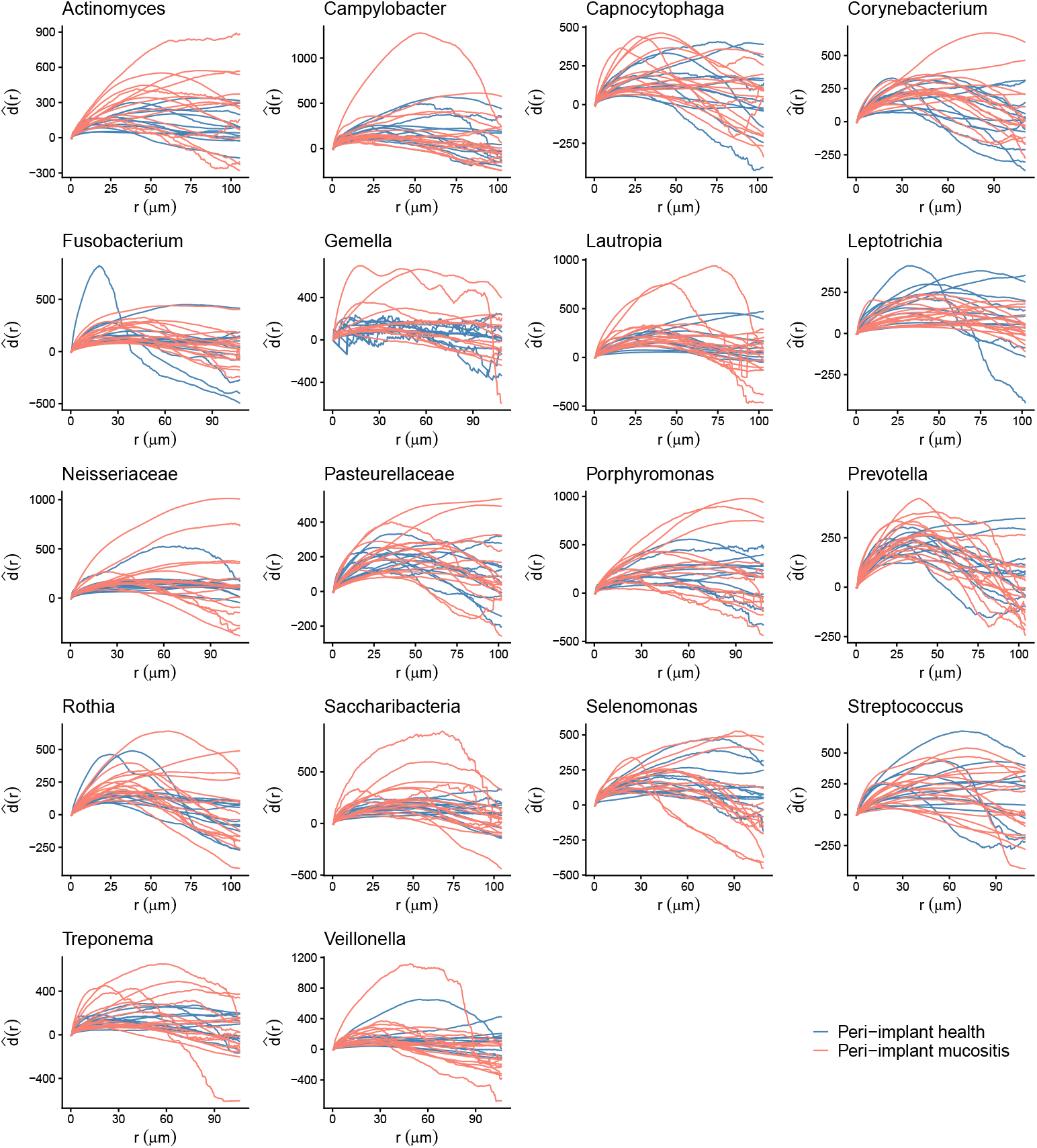
Estimated *d*(*r*) curves for each microbial genus in the oral biofilm dataset, with mucositis images colored red and healthy images colored blue.

**Fig. 7.**
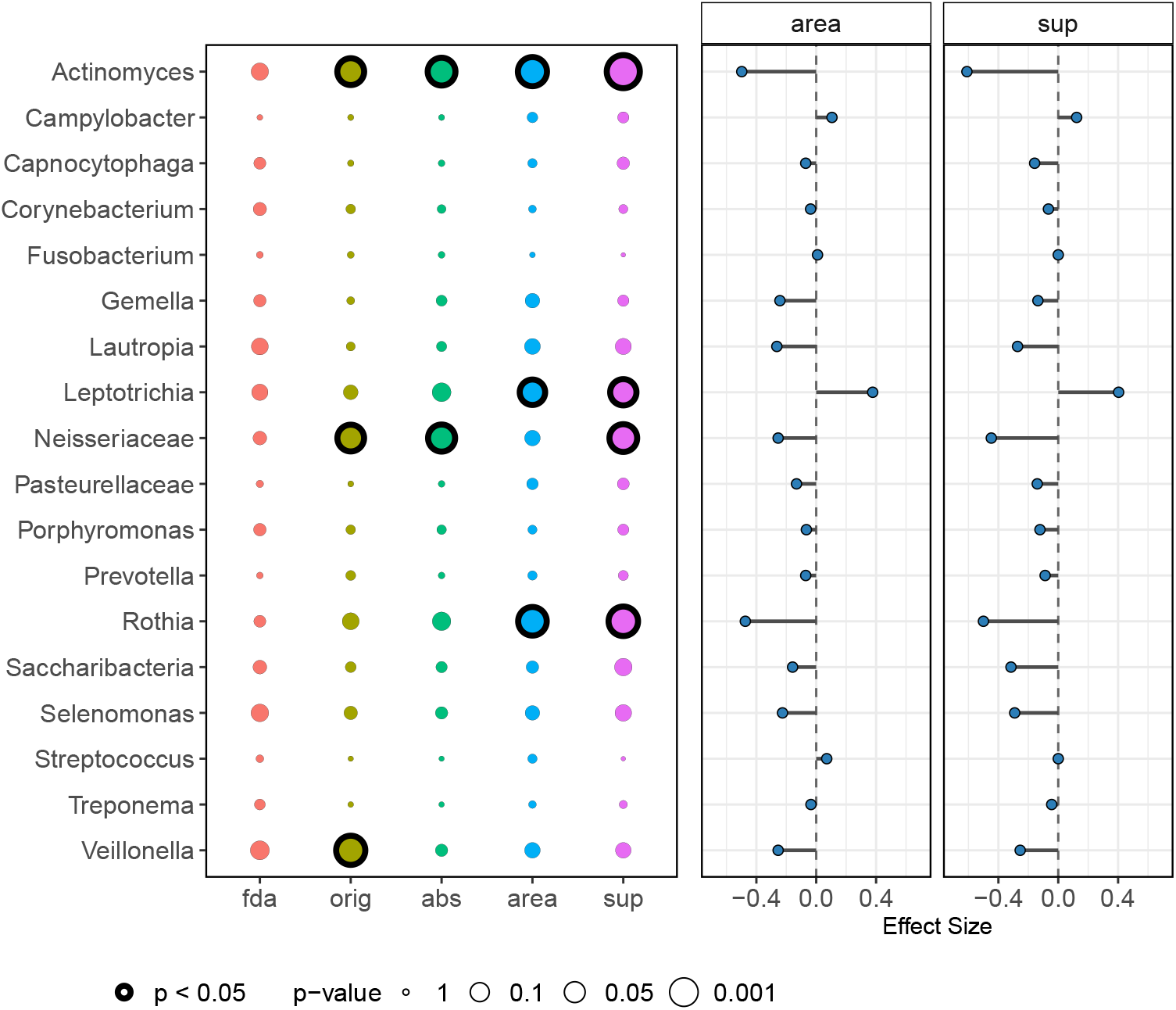
Application to the oral biofilm dataset. The left panel shows taxon-specific *p*-values for all MiSA tests and the competing method, fda. Significant results (*p <* 0.05) are highlighted with bordered dots. The right panel shows effect sizes for MiSA-area and MiSA-sup.

At a significance level of *α* = 0.05, MiSA identified several microbial genera with spatial differences between the two diagnostic groups, which may contribute to spatial dysbiosis, whereas fda detected none.

*Actinomyces* showed spatial differences in all four MiSA tests. MiSA-orig (*p* = 0.0260) and MiSA-abs (*p* = 0.0168) indicated that its spatial pattern differs between groups at specific radii. The difference was also evident in the most striking spatial patterns, captured by MiSA-sup (*p* = 0.0009, effect size = ™0.610), and in the integrated spatial signal, captured by MiSA-area (*p* = 0.0087, effect size = ™0.498). The negative effect sizes quantitatively reflect larger a deviation from CSR in the mucosi-tis group, pointing to a more pronounced spatial pattern than in the healthy group.

Consistent with these results, the 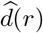 curves of both groups lie mostly above 0, but the mucositis group deviates further from CSR across radii (Figure 6). Together, these results suggest that *Actinomyces* forms stronger clusters in mucositis biofilms, whereas in healthy biofilms its spatial distribution is less structured and closer to CSR. These findings are consistent with the representative images in Figure S1 (Additional file 1), where *Actinomyces* forms small colonies scattered throughout the peri-implant health images, but fewer, larger patches with sharp boundaries in the peri-implant mucositis images (stronger clustering). Existing literature also indicates that *Actinomyces* tends to adopt more structured forms under disease conditions, providing further support for our findings. *Actinomyces* is a common oral commensal that can become pathogenic when oral homeostasis is disrupted [22]. *Actinomyces* species also contribute to oral biofilm formation and dental plaque-related diseases [23], and dental plaque is a spatially structured community whose organization differs between health and disease [24]. More generally, bacteria in oral biofilms, including *Actinomyces*, are spatially organized and embedded in an extracellular polymeric matrix that provides structural stability and protection against antimicrobials [25].

*Leptotrichia* exhibits a potential contribution to spatial dysbiosis based on MiSA-area (*p* = 0.0473) and MiSA-sup (*p* = 0.0331), suggesting that its spatial differences between groups are most detectable through global summaries 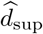 and 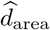. In contrast to *Actinomyces, Leptotrichia* showed medium positive effect s izes (0.377 for MiSA-area and 0.404 for MiSA-sup; Table 2), indicating more pronounced spatial structure in the healthy group than in the mucositis group. This is illustrated in Figure 6, where the *d*(*r*) curves of the healthy group generally lie farther from CSR. Studies of *Leptotrichia buccalis* reveal that it lacks common pili (fimbriae) and an S-layer, which are essential for building rigid, multi-layered physical structures. Instead, it has a “mosaic” of surface ridges that may be less effective at maintaining complex 3D architecture during active disease [26, 27]. Other taxa that may contribute to spatial dysbiosis identified by MiSA include *Neisseriaceae, Rothia*, and *Veillonella*, with details provided in Additional file 1: Section S2.

Overall spatial dysbiosis of the entire microbial community was detected by MiSA-sup, aggregating *p*-values across the 18 taxa (combined *p* = 0.0148), whereas the combined *p*-values for the other MiSA tests were not significant (MiSA-orig: *p* = 0.1295; MiSA-abs: *p* = 0.2208; MiSA-area: *p* = 0.1028). This is because MiSA-sup focuses on the largest deviation from CSR, thus is sensitive to strong but scale-localized signals, which are weakened when deviations are aggregated over radii. The detected spatial dysbiosis is consistent with previous studies showing that the spatial organization of oral biofilms is altered during disease progression. In dental caries, bacteria organize into rotund-shaped structures with a core of *Streptococcus mutans* surrounded by layers of other species [28], and the extracellular matrix of such mixed-species biofilms protects bacteria from host immune responses and antimicrobials [29].

## 4 Discussion

The emerging image-based spatial microbiome data provide additional spatial resolution to study differences in microbial spatial organization between health and disease, beyond traditional compositional comparisons. These data have unique characteristics that are not present in the relatively well-studied spatial transcriptomics and proteomics data. First, tens of taxa are of interest altogether, which reduces the power of methods rooted in pairwise co-occurrence of cell types and limits methods based on spatial entropy, which depends only on the proportions of taxa and not on their identities, so individual taxa’s spatial aberrations are obscured. Second, microbiome spatial patterns are heterogeneous even within diagnostic groups, reducing the power of functional data-based methods, which rely on systematic shifts of curves between groups. Third, bacteria naturally form small colonies and thus show clustering at small scales even in well-mixed communities, so methods that test each image against CSR are driven by this natural clustering rather than by between-group differences.

To effectively analyze these data, we proposed MiSA, the first hypothesis testing framework for identifying spatial dysbiosis in image-based microbiome data. By comparing each taxon’s deviation from CSR, summarized by *d*(*r*) across radii, between clinical groups. MiSA focuses on differences in spatial organization rather than on clustering shared by all images, such as natural bacterial colonies. It also accommodates many taxa simultaneously, highly heterogeneous spatial patterns within groups, and signals that vary across spatial scales in both magnitude and direction. The four MiSA tests are complementary and can be chosen according to the specific scientific question. MiSA-orig and MiSA-abs are suited to applications in which specific spatial scales are of primary interest, with MiSA-orig also accounting for the direction of deviation (clustering or regularity). MiSA-sup and MiSA-area are suited to assessing spatial structure over the whole image, with MiSA-sup sensitive to strong signals at any scale and MiSA-area accumulating signals across scales. When no specific spatial scale or direction of deviation is hypothesized, we recommend MiSA-sup as the default, as it does not assume that signals occur at particular radii and is sensitive to strong, scale-localized signals.

In real data-based simulation studies, MiSA controlled the type I error and achieved substantially higher power than the competing method borrowed from spatial proteomics, fda, at both the individual-taxon and community levels. Applied to the HiPR-FISH oral biofilm data, exploratory taxon-level analyses by MiSA suggested that five taxa, *Actinomyces, Leptotrichia, Neisseriaceae, Rothia*, and *Veillonella*, had spatial patterns that differed between peri-implant health and peri-implant mucositis and may contribute to spatial alterations of the microbiome. At the community level, MiSA-sup detected spatial dysbiosis. None of these signals was detected by fda, because fda primarily detects systematic shifts in the spatial summary curves, which are weak when spatial patterns vary widely within groups. As MiSA requires only the locations and taxonomic identities of individual cells, it is applicable to other image-based microbiome mapping technologies beyond HiPR-FISH, such as CLASI-FISH [30] and SEER-FISH [31].

MiSA has several limitations. First, MiSA is implemented following microbial segmentation and taxonomic classification, and therefore relies on sufficient accuracy in these steps to ensure valid biological inference. Future extensions could explore segmentation-free approaches to reduce the impact of segmentation errors. Second, because spatial signals may arise at any spatial scale, results may depend on the choice of MiSA variant and radii *r*, which investigators should therefore select a priori according to the specific biological question.

## 5 Conclusions

We present MiSA, the first statistical framework for identifying disease-associated spatial dysbiosis in image-based microbiome data. MiSA compares each taxon’s deviation from CSR between clinical groups, at specific radii or through global summaries, capturing signals that vary across spatial scales. Combining these taxon-level results then allows spatial dysbiosis to be assessed at the community level. In simulations and in an application to a real oral biofilm study, MiSA detected spatial dysbiosis that a competing method borrowed from spatial proteomics failed to detect. MiSA therefore opens an avenue for powerful and robust analysis of microbiome spatial dysbiosis as such image-based spatial microbiome data become increasingly available.

## List of abbreviations

HiPR-FISH: (High Phylogenetic Resolution Fluorescence in situ Hybridization)
MiSA: (Microbiome Spatial Aberration)
CSR: (Complete Spatial Randomness)
fda: (functional data analysis)
HMP: (harmonic mean *p*-value)

## Declarations

### Ethics approval and consent to participate

Not applicable. Because this study only involved secondary analysis of an existing, de-identified dataset, it is not considered human subject research.

### Consent for publication

Not applicable.

### Availability of data and material

The segmented oral biofilm spatial microbiome data analysed in this work are available on Zenodo (https://doi.org/10.5281/zenodo.24094726); the original data were described in [6]. The R package `MiSA` is available at https://github.com/liuruoqian/MiSA in formats appropriate for Macintosh or Windows systems. A vignette demonstrating use of the package is included (can be accessed at https://liuruoqian.github.io/MiSA/articles/MiSA_vignette.html).

## Authors’ contributions

R.L. developed the method, analyzed the data, and wrote the manuscript. B.Gr. and generated the data, and contributed to the conception, methodological developments, and presentation. S.S., D.D., A.R., Y.D., B.Gu. contributed to methodological developments and presentation. N.Z. contributed to the conception, methodological developments, and reviewed the manuscript. W.L. conceived the study and critically reviewed the manuscript.

## Acknowledgements

Not applicable.

## Endnotes

Not applicable.

## Additional files

**Additional file 1** (.pdf) — Supplementary information. Section S1: Data generation for the illustrative example in Figure 2. Section S2: Additional taxa that may contribute to spatial dysbiosis in the oral biofilm study. Figure S1: Additional figure for the oral biofilm data analysis.

## References

[1] DeGruttola, A.K., Low, D., Mizoguchi, A., Mizoguchi, E.: Current understanding of dysbiosis in disease in human and animal models. Inflammatory bowel diseases 22(5), 1137–1150 (2016)

[2] Önata, E.T., Özdemir, Ö.: Microbiome, dysbiosis and use of probiotics in various diseases. World Journal of Virology 14(2), 99574 (2025)

[3] Hou, K., Wu, Z.-X., Chen, X.-Y., Wang, J.-Q., Zhang, D., Xiao, C., Zhu, D., Koya, J.B., Wei, L., Li, J., et al.: Microbiota in health and diseases. Signal transduction and targeted therapy 7(1), 135 (2022)

[4] Sabih Ur Rehman, S., Nasar, M.I., Mesquita, C.S., Al Khodor, S., Notebaart, R.A., Ott, S., Mundra, S., Arasardanam, R.P., Muhammad, K., Alam, M.T.: Integrative systems biology approaches for analyzing microbiome dysbiosis and species interactions. Briefings in Bioinformatics 26(4), 323 (2025)

[5] Shi, H., Shi, Q., Grodner, B., Lenz, J.S., Zipfel, W.R., Brito, I.L., De Vlaminck, I.: Highly multiplexed spatial mapping of microbial communities. Nature 588(7839), 676–681 (2020)

[6] Grodner, B., Wu, D.T., Hahm, S., Takayasu, L., Wen, N., Kim, D.M., Chen, C.-Y., De Vlaminck, I.: Microscale spatial dysbiosis in oral biofilms associated with disease. bioRxiv, 2024–07 (2024)

[7] Hajishengallis, G., Lamont, R.J., Koo, H.: Oral polymicrobial communities: Assembly, function, and impact on diseases. Cell host & microbe 31(4), 528–538 (2023)

[8] Kim, D., Koo, H.: Spatial design of polymicrobial oral biofilm in its native disease state. Journal of Dental Research 99(6), 597–603 (2020)

[9] Greve, D., Moter, A., Kleinschmidt, M.C., Pfäfflin, F., Stegemann, M.S., Kursawe, L., Grubitzsch, H., Falk, V., Kikhney, J.: Rothia aeria and rothia dentocariosa as biofilm builders in infective endocarditis. International Journal of Medical Microbiology 311(2), 151478 (2021)

[10] Canete, N.P., Iyengar, S.S., Ormerod, J.T., Baharlou, H., Harman, A.N., Patrick, E.: spicyr: spatial analysis of in situ cytometry data in r. Bioinformatics 38(11), 3099–3105 (2022)

[11] Seal, S., Neelon, B., Angel, P.M., O’Quinn, E.C., Hill, E., Vu, T., Ghosh, D., Mehta, A.S., Wallace, K., Alekseyenko, A.V.: Spaceanova: Spatial co-occurrence analysis of cell types in multiplex imaging data using point process and functional anova. Journal of Proteome Research 23(4), 1131–1143 (2024)

[12] Masotti, M., Osher, N., Eliason, J., Rao, A., Baladandayuthapani, V.: Dimple: an r package to quantify, visualize, and model spatial cellular interactions from multiplex imaging with distance matrices. Patterns 4(12) (2023)

[13] Vu, T., Seal, S., Ghosh, T., Ahmadian, M., Wrobel, J., Ghosh, D.: Funspace: a functional and spatial analytic approach to cell imaging data using entropy measures. PLOS Computational Biology 19(9), 1011490 (2023)

[14] Wrobel, J., Soupir, A.C., Hayes, M.T., Peres, L.C., Vu, T., Leroux, A., Fridley, B.L.: mxfda: a comprehensive toolkit for functional data analysis of single-cell spatial data. Bioinformatics advances 4(1), 155 (2024)

[15] Ripley, B.D.: The second-order analysis of stationary point processes. Journal of applied probability 13(2), 255–266 (1976)

[16] Ripley, B.D.: Modelling spatial patterns. Journal of the Royal Statistical Society: Series B (Methodological) 39(2), 172–192 (1977)

[17] Baddely, A., Rubak, E., Turner, R.: Spatial point patterns: methodology and applications with r. Chapman and Hall/CRC Press, London (2015)

[18] Matérn, B.: Spatial variation: Meddelanden fran statens skogsforskningsinstitut. Lecture Notes in Statistics 36, 21 (1960)

[19] Wilson, D.J.: The harmonic mean p-value for combining dependent tests. Proceedings of the National Academy of Sciences 116(4), 1195–1200 (2019)

[20] Peres, F.F.: Effect sizes for nonparametric tests. Biochemia Medica 36(1), 010101 (2025)

[21] Pateiro-López, B., Rodríguez-Casal, A.: Generalizing the convex hull of a sample: the r package alphahull. Journal of Statistical software 34, 1–28 (2010)

[22] Thukral, R., Shrivastav, K., Mathur, V., Barodiya, A., Shrivastav, S.: Actinomyces: a deceptive infection of oral cavity. Journal of the Korean Association of Oral and Maxillofacial Surgeons 43(4), 282 (2017)

[23] Akbar, P., Salahuddin, N., Ahmad, Z., Shah, S.Z., Shah, F., Maknoon, D.: Role of actinomyces species in oral biofilm formation and dental plaque-related diseases. Innovative Research in Applied, Biological and Chemical Sciences 2(1), 120–125 (2024)

[24] Borisy, G.G., Valm, A.M.: Spatial scale in analysis of the dental plaque micro-biome. Periodontology 2000 86(1), 97–112 (2021)

[25] Gerardi, D., Bernardi, S., Bruni, A., Falisi, G., Botticelli, G.: Characterization and morphological methods for oral biofilm visualization: where are we nowadays? AIMS microbiology 10(2), 391 (2024)

[26] Eribe, E.R.K., Olsen, I.: Leptotrichia species in human infections. Anaerobe 14(3), 131–137 (2008)

[27] Smith, S., Murray, R., Hall, M.: The surface structure of leptotrichia buccalis. Canadian journal of microbiology 40(2), 90–98 (1994)

[28] Kim, D., Barraza, J.P., Arthur, R.A., Hara, A., Lewis, K., Liu, Y., Scisci, E.L., Hajishengallis, E., Whiteley, M., Koo, H.: Spatial mapping of polymicrobial communities reveals a precise biogeography associated with human dental caries. Proceedings of the National Academy of Sciences 117(22), 12375–12386 (2020)

[29] Kurtzman, G.M., Horowitz, R.A., Johnson, R., Prestiano, R.A., Klein, B.I.: The systemic oral health connection: Biofilms. Medicine 101(46), 30517 (2022)

[30] Valm, A.M., Welch, J.L.M., Borisy, G.G.: Clasi-fish: principles of combinatorial labeling and spectral imaging. Systematic and applied microbiology 35(8), 496–502 (2012)

[31] Cao, Z., Zuo, W., Wang, L., Chen, J., Qu, Z., Jin, F., Dai, L.: Spatial profiling of microbial communities by sequential fish with error-robust encoding. Nature Communications 14(1), 1477 (2023)

