## Supplementary Materials for "Identifying disease-associated dysbiosis in image-based spatial microbiome via MiSA"

**Additional File 1**

This additional file contains (1) a description of how the data for the illustrative example in the main text (Figure 2) were generated, (2) details for additional taxa that may contribute to spatial dysbiosis in the oral biofilm study, and (3) an additional figure for the oral biofilm data analysis.

### S1 Data generation for Figure 2 in the main text

Point Pattern 1 is formed by first generating a uniform Poisson point process of "parent" points with intensity `kappa = 25`. Each "parent" point is then replaced by a random cluster of "offspring" points, where the number of points per cluster follows a Poisson distribution with mean `mu = 8`. The "offspring" points distributed uniformly within a disc of radius `scale = 0.03` centered at the "parent" point. In contrast, Point Pattern 2 reflects the spatial expansion of microbial colonies. Specifically, clusters have a radius of 0.1, contain an average of 40 "offspring" points, and arise from "parent" points with a lower intensity, `kappa = 5`. Consequently, Pattern 2 consists of fewer but larger clusters. Both patterns have the same overall intensity, `kappa * mu = 200`.

### S2 Additional taxa potentially contributing to spatial dysbiosis in oral biofilms

*Neisseriaceae* showed spatial differences under MiSA-orig ($p = 0.0273$), MiSA-abs ($p = 0.0233$), and MiSA-sup ($p = 0.0172$, effect size $= -0.447$), indicating differences at specific radii as well as in the maximum deviation from CSR. The negative effect size indicates a more strongly clustered arrangement of *Neisseriaceae* in the mucositis group. Consistently, the $\widehat{d}(r)$ curves of the mucositis group lie farther from CSR than those of the healthy group (Figure 6), and *Neisseriaceae* appears more clustered in the representative mucositis images (Figure S1). Several *Neisseriaceae* species, including *Kingella oralis*, *Neisseria elongata*, and *Neisseria mucosa*, are specialized to dental plaque, with nitrate reduction and denitrification capabilities that may favor persistence within structured biofilms [3]. Because plaque organization differs between health and disease [1, 6], the stronger clustering of *Neisseriaceae* in mucositis may reflect a disease-associated shift in plaque organization.

*Rothia* showed spatial differences under MiSA-area ($p = 0.0113$) and MiSA-sup ($p = 0.0071$), with medium negative effect sizes ($-0.474$ and $-0.500$, respectively), indicating that its spatial arrangement deviates more from CSR in the mucositis group. Consistently, *Rothia* occupies large, contiguous patches in the mucositis images (Figure S1, particularly panel d), whereas in the healthy images it forms smaller clusters scattered throughout the image. *Rothia* is a recognized biofilm former that colonizes oral surfaces and dental plaque [7], and it can form structured biofilms in infection, as in infective endocarditis [4].

*Veillonella* showed spatial differences under MiSA-orig ($p = 0.0085$), indicating differences at specific radii when the direction of deviation from CSR is taken into account. As shown in Figure 6, the $\widehat{d}(r)$ curves of several mucositis images are positive at small $r$ but become negative at larger $r$, whereas most healthy-group curves remain close to 0. This indicates that in mucositis, *Veillonella* forms small clusters that are themselves more regularly spaced than expected under CSR, while in health it is distributed more randomly. This pattern is also shown in Figure S1, where *Veillonella* is spread across the healthy images but forms distinct clusters in the mucositis images. These findings are consistent with the existing literature. *Veillonella* is a bridging taxon that shapes the architecture of multi-species oral biofilms and can act as a pathobiont that alters biofilm spatial structure in oral diseases [5, 9, 2, 8].

### S3 Supplementary Figure

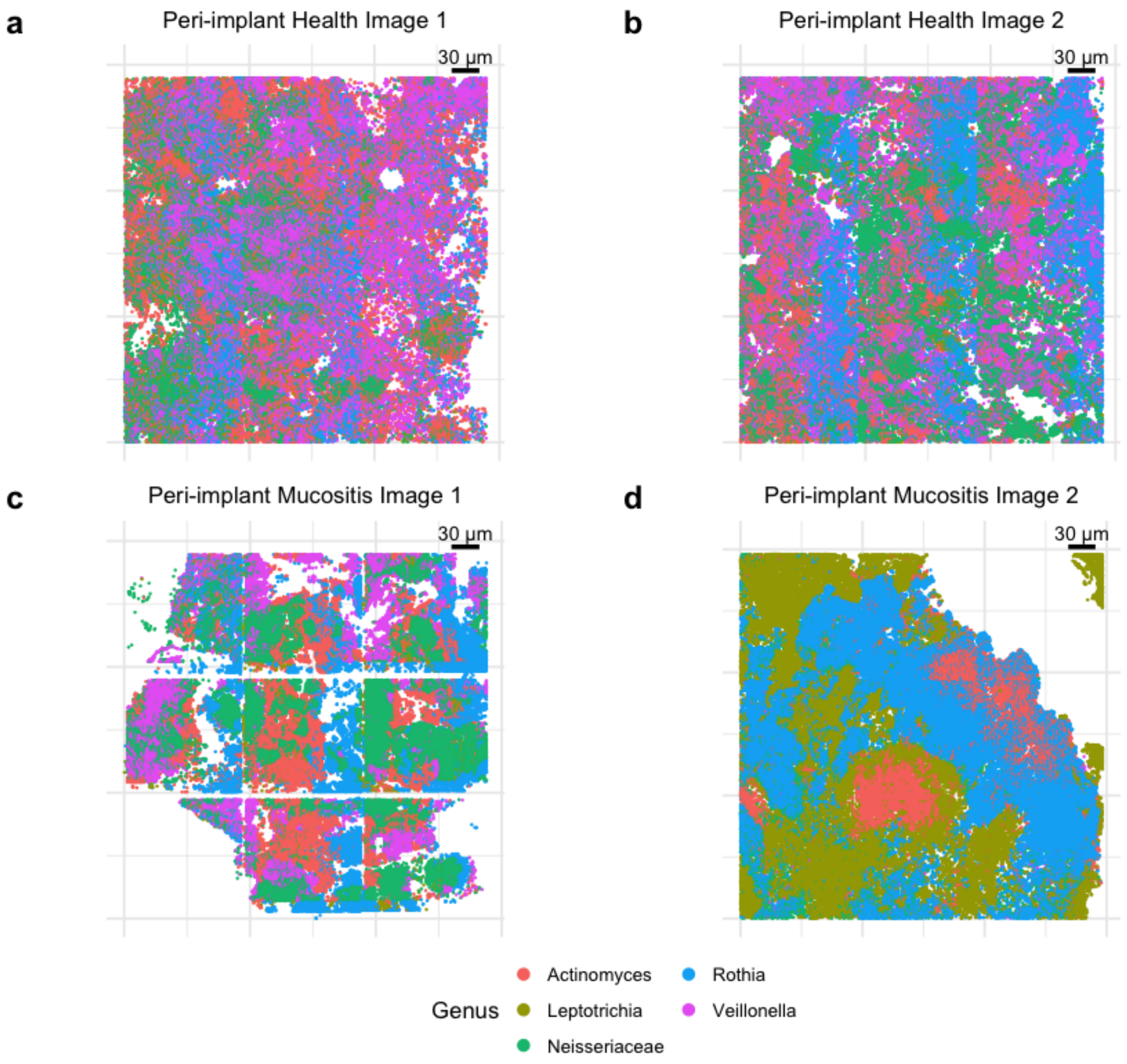


**Figure S1:** Representative examples of segmented oral biofilm spatial microbiome data, showing the genus identities of taxa that may contribute to spatial dysbiosis.

### References


[1] Gary G Borisy and Alex M Valm. Spatial scale in analysis of the dental plaque microbiome. *Periodontology 2000*, 86(1):97–112, 2021.

[2] Louis Dorison, Nathalie Béchon, Camille Martin-Gallausiaux, Susan Chamorro-Rodriguez, Yakov Vitrenko, Rania Ouazahrou, Romain Villa, Julien Deschamps, Romain Briandet, Simonetta Gribaldo, et al. Identification of veillonella parvula and streptococcus gordonii adhesins mediating co-aggregation and its impact on physiology and mixed biofilm structure. *MBio*, 15(12):e02171–24, 2024.

[3] Jonathan J Giacomini, Julian Torres-Morales, Floyd E Dewhirst, Gary G Borisy, and Jessica L Mark Welch. Spatial ecology of the neisseriaceae family in the human oral cavity. *Microbiology Spectrum*, 13(5):e03275–24, 2025.

[4] Dustin Greve, Annette Moter, Malte Christian Kleinschmidt, Frieder Pfäfflin, Miriam Songa Stegemann, Laura Kursawe, Herko Grubitzsch, Volkmar Falk, and Judith Kikhney. Rothia aeria and rothia dentocariosa as biofilm builders in infective endocarditis. *International Journal of Medical Microbiology*, 311(2):151478, 2021.

[5] Junli Li, Hejing Wang, Na Li, Yikui Zhang, Xin Lü, and Bianfang Liu. Antibiotic susceptibility and biofilm-forming ability of veillonella strains. *Anaerobe*, 78:102667, 2022.

[6] Allan Radaic and Yvonne L Kapila. The oralome and its dysbiosis: New insights into oral microbiome-host interactions. *Computational and structural biotechnology journal*, 19:1335–1360, 2021.

[7] Syahdiana Waty and Nurul Hidayah. Identification of bacteria in dental plaque. *Contagion: Scientific Periodical Journal of Public Health and Coastal Health*, 7(1):1–10, 2025.

[8] Yuan Wei, Yu Zhang, Yuan Zhuang, Yifei Tang, Hua Nie, Yequan Haung, Ting Liu, Weidong Yang, Fuhua Yan, and Yanan Zhu. Veillonella parvula acts as a pathobiont promoting the biofilm virulence and cariogenicity of streptococcus mutans in adult severe caries. *Microbiology Spectrum*, 12(11):e04318–23, 2024.

[9] Peng Zhou, Daniel Manoil, Georgios N Belibasakis, and Georgios A Kotsakis. Veillonellae: beyond bridging species in oral biofilm ecology. *Frontiers in oral health*, 2:774115, 2021.
